# On the evolution of the plant phytochrome chromophore biosynthesis

**DOI:** 10.1101/2022.08.22.504806

**Authors:** Federica Frascogna, Benjamin Ledermann, Jana Hartmann, Eugenio Pérez Patallo, Fjoralba Zeqiri, Eckhard Hofmann, Nicole Frankenberg-Dinkel

## Abstract

Phytochromes are biliprotein photoreceptors present in plants, algae, certain bacteria and fungi. Land plant phytochromes use phytochromobilin (PΦB) as the bilin chromophore. Phytochromes of streptophyte algae, the clade within which land plants evolved, employ phycocyanobilin (PCB), leading to a more blue-shifted absorption spectrum. Both chromophores are synthesized by ferredoxin-dependent bilin reductases (FDBRs) starting from biliverdin IXα (BV). In cyanobacteria and chlorophyta, BV is reduced to PCB by the FDBR phycocyanobilin:ferredoxin oxidoreductase (PcyA), whereas, in land plants, BV is reduced to PФB by phytochromobilin synthase (HY2). However, phylogenetic studies suggested the absence of any ortholog of PcyA in streptophyte algae and the presence of only PФB biosynthesis related genes (*HY2*). The HY2 of the early diverging streptophyte alga *Klebsormidium nitens* (formerly *Klebsormidium flaccidum*) was already indirectly indicated to be involved in PCB biosynthesis. Here, we overexpressed and purified a His_6_-tagged variant of *K. nitens* HY2 (KflaHY2) in *E. coli*. Employing anaerobic bilin reductase activity assays and coupled phytochrome assembly assays, we were able to confirm the product and to identify intermediates of the reaction. Site-directed mutagenesis revealed two aspartate residues critical for catalysis. While it was not possible to convert KflaHY2 into a PΦB-producing enzyme by simply exchanging the catalytic pair, the biochemical investigation of two additional members of the HY2 lineage enabled us to define two distinct clades, the PCB-HY2 and the PΦB-HY2 clade. Overall, our study gives insight into the evolution of the HY2 lineage of FDBRs.

## Introduction

Photosynthesis is arguably one of the most significant biological processes on earth. To harvest light efficiently, photosynthetic organisms possess chlorophyll light-harvesting antennae incorporated in their photosystems. Moreover, some of these organisms are also capable to sense the quality and intensity of the harvested light. To do that, they employ several types of photoreceptors, which are able to absorb photons and to modulate biological activities based on this information (Möglich et al., 2010). Among these photoreceptors, phytochromes can be found in plants, algae, certain bacteria and fungi (Sharrock, 2008). Phytochromes are capable to perceive red and far-red light, by employing open-chain tetrapyrroles as light-sensing chromophores. Although phylogenetically closely related, land plants and streptophyte algae employ different bilins in their phytochromes. Indeed, as far as is known, land plants use the dark green pigment phytochromobilin (PΦB), whereas streptophyte algae employ the cyan pigment phycocyanobilin (PCB) (Wu et al., 1997). The transition from PCB to PΦB as the phytochrome chromophore appears to have coincided with the origin of land plants (Rockwell et al., 2017). The different bound chromophore leads to a blue shift of the photocycles in the phytochromes of streptophyte algae compared to the ones of land plants.

The light-sensing open-chain tetrapyrroles employed in the photoreceptors are all derived from heme. The first step in the biosynthesis of the bilins is the cleavage of the heme macrocycle at the α-methine bridge mediated by heme oxygenases (HOs). This reaction yields the open-chain tetrapyrrole biliverdin IXα (BV), liberating H_2_O, CO and Fe^2+^ (Cornejo and Beale, 1988). BV is then subsequently reduced to the specific bilins by a class of enzymes called ferredoxin-dependent bilin reductases (FDBRs) (Figure S1) (Frankenberg et al., 2001; Frankenberg and Lagarias, 2003). In land plants, the enzyme phytochromobilin synthase (HY2) reduces BV in a two-electron reduction at the A-ring 2,3,3^1^,3^2^-diene system to the dark green pigment PΦB (Figure 1) (Kohchi et al., 2001; McDowell and Lagarias, 2001). In cyanobacteria and chlorophyta, phycocyanobilin:ferredoxin oxidoreductase (PcyA) catalyzes the four-electron reduction of BV to the cyan pigment PCB via the intermediate 18^1^,18^2^-dihydrobiliverdin (18^1^,18^2^-DHBV) (Figure 1) (Frankenberg and Lagarias, 2003). Interestingly, the streptophyte alga *Klebsormidium nitens* is known to employ PCB in its phytochromes but only possesses a HY2 homolog (Rockwell and Lagarias, 2017). Using coexpression studies, it was suggested that the HY2 of *K. nitens* (named KflaHY2) is able to synthesize PCB and not PΦB, like its higher plant homolog (Rockwell et al., 2017). Here we provide direct biochemical evidence that recombinant KflaHY2 is able to reduce BV to PCB. In addition, we investigated the reaction in more detail, identifying important catalytic residues. Biochemical characterization of two additional members of the HY2 lineage furthermore enabled us to define two distinct clades.

**Figure 1.**
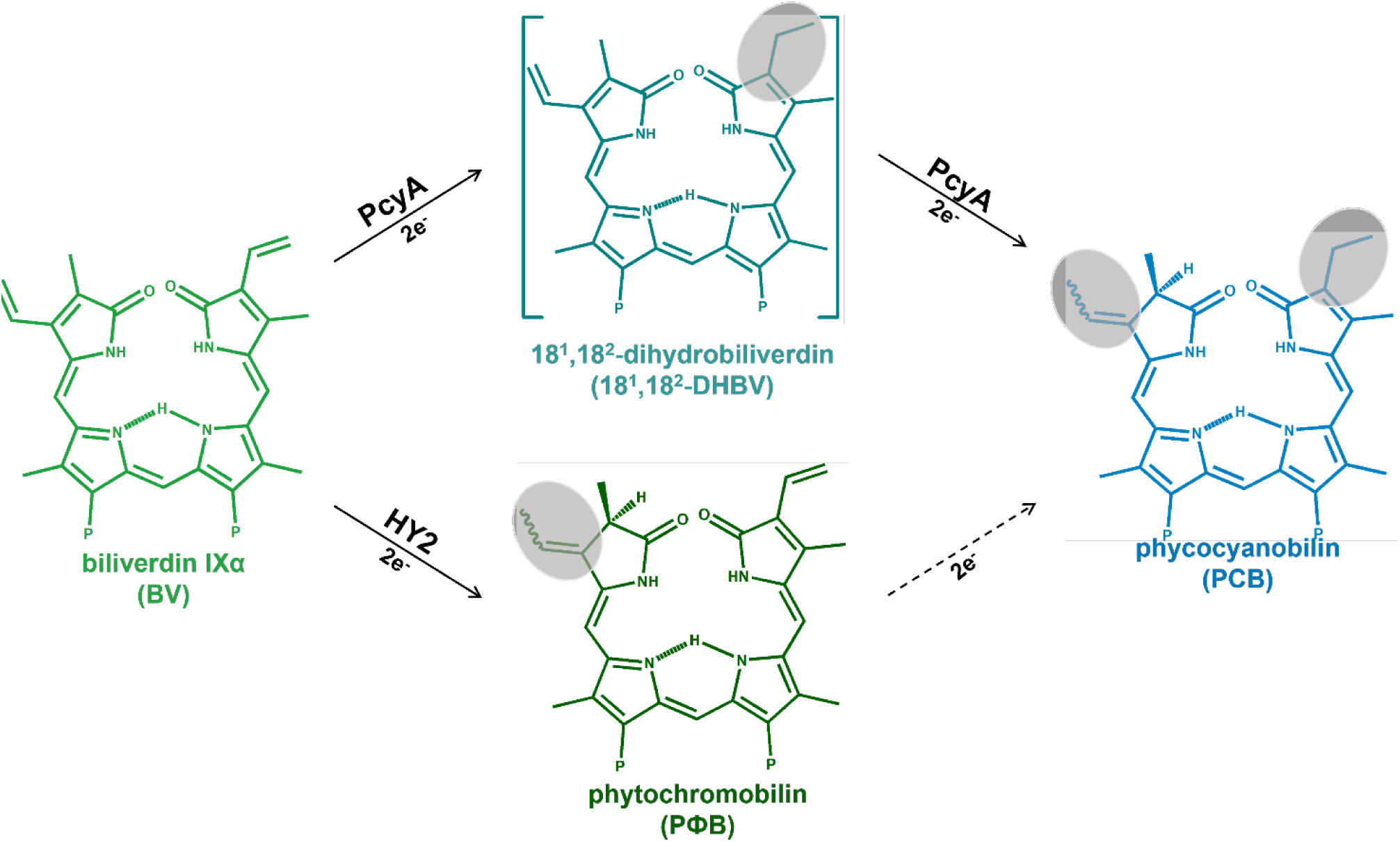
Overview of the reactions catalyzed by the FDBRs HY2 and PcyA. BV is the substrate of the FDBRs in question. “P” indicates the propionate side chains. Land Plants HY2 catalyzes the two-electron reduction of BV to PΦB. PcyA catalyzes the four-electron reduction of BV to PCB via the intermediate 18^1^,18^2^-DHBV. Dashed line represents a possible second pathway to PCB.

Overall, our data give more insight into the evolution of this interesting enzyme family and on the evolution of plants phytochromes.

## Materials and Methods

### Chemicals

All chemicals used in this study were ACS grade or better. Most of the components for the anaerobic bilin reductase activity tests were also purchased from Sigma-Aldrich (St. Louis, Missouri, USA), except for biliverdin IXα, which was purchased from Frontier Scientific (Logan, Utah, USA). Ferredoxin (PetF) and FNR (PetH) were recombinantly expressed and purified as described elsewhere (Dammeyer et al., 2008; Busch et al., 2011). HPLC-grade acetone, formic acid and acetonitrile were purchased from VWR Chemicals (Radnor, Pennsylvania, USA).

### Plasmids

The plasmid pET28a_*KflaHY2* was gifted by Prof. J. Clark Lagarias (UC Davis, California, USA). Site-directed mutants of KflaHY2 were generated in the pET28a_*KflaHY2* plasmid using the QuikChange^®^ Lightning kit (Agilent, Santa Clara, California, USA) employing primers listed in Table S1. The variant proteins were produced and purified according to the protocol used for KflaHY2 unless stated otherwise. The plasmids pGEX-4T-1_*CepuHY2* and pGEX-4T-1_*NediHY2* were purchased as already cloned by GenScript (Piscataway, New Jersey, USA). The full list of employed constructs can be found in Supplemental Materials (Table S2).

### Production and purification of recombinant proteins

For the production of recombinant KflaHY2 and variants, 2 L of LB medium containing the appropriate antibiotics was inoculated 1:100 with an overnight culture of *E. coli* BL21(DE3) carrying the respective plasmids. Cells were grown at 37°C and 100 rpm (Innova^®^ 44, New Brunswick Scientific, Enfield, Connecticut, USA) to an OD_600_ of 0.4-0.6. The temperature was decreased to 17°C and gene expression was induced by the addition of isopropyl-β-D-thiogalactopyranoside (IPTG) to a final concentration of 1 mM (for pET28a_*KflaHY2*) or 0.5 mM (for pET28a_*KflaHY2*_N105D, pET28a_*KflaHY2*_D122N, pET28a_*KflaHY2*_D242N, pET28a_*KflaHY2*_N105D_D122N, pET28a_*KflaHY2*_N105D_D242N). The cultures were incubated under shaking for additional 19 h and harvested at 4°C by centrifugation for 10 min at 17000 g (Sorvall^TM^ LYNX^TM^ 6000, Rotor F9). The pellets were resuspended in “binding buffer” (20 mM sodium phosphate pH 7.4; 500 mM NaCl). After the addition of a spatula tip of DNaseI (Applichem, Darmstadt, Germany) and lysozyme (Sigma-Aldrich), the suspension was kept on ice for 30 min. The cells were disrupted by sonication for 5 min (Bandelin Sonopuls HD 2200; tip KE76; 5’’ pulses, 10’’ pauses; cycle 6/10; ≈ 40% power output) and centrifuged for 45 min at 4°C and 50000 g (Sorvall^TM^ LYNX^TM^ 6000, rotor T29-8). The crude extract was loaded onto a gravity flow column containing 2 mL of TALON® Superflow™ resin (Cytiva, Marlborough, Massachusetts, USA). The column was washed with 10 column volumes (CV) of binding buffer and the elution was performed using 4 CV of “elution buffer” (20 mM sodium phosphate pH 7.4; 500 mM NaCl; 500 mM imidazole). Protein containing fractions were pooled and dialyzed overnight against TES-KCl buffer (25 mM TES/KOH pH 7.5; 100 mM KCl; 10% glycerol).

Production of NediHY2 and CepuHY2 was carried out adding minor adjustments to the above-mentioned protocol. During expression, after an OD_600_ of 0.4-0.6 was reached, the main culture was subjected to a cold shock at 4°C for 1h, before inducing with 0.5 mM IPTG. The cultures were incubated under shaking at 17°C for additional 19 h before harvesting. The pellets were resuspended in “lysis buffer” (50 mM Tris-HCl pH 7.5; 150 mM NaCl) and a spatula tip of DNAse was added to the resuspension alongside 1 mM PMSF. After homogenizing the resuspension with a douncer, lysis was carried out using a Microfluidics^TM^ M-110L microfluidizer (3-5 cycles, 800-1000 bar). The disrupted cells were subjected to ultracentrifugation at 4°C and 235000 g for 1h (Beckman Optima™ L-80 XP, rotor Ti45). The resulting supernatant was loaded onto an Äkta purifier system (GE HealthCare, Chicago, Illinois, USA) equipped with a 25 mL self-packed 20/26 GSTrap column (GE HealthCare). A linear gradient containing 50 mM reduced glutathione in “lysis buffer” was used for elution. Protein containing fractions were pooled and buffer exchange was carried out using a PD-10 column (GE Healthcare) using TES-KCl buffer (25 mM TES/KOH pH 7.5; 100 mM KCl; 10% glycerol). The fusion protein was cleaved using Thrombin (from bovine plasma, 50 NIH units/mg dry wt) following the manufacturer’s instructions (Sigma-Aldrich). The following day, after concentrating the protein (Amicon® Ultra MWCO 30 kDa, Merck, Darmstadt, Germany), a second GST-tag purification was performed and the peak fractions were pooled and concentrated (Amicon® Ultra MWCO 10 kDa, Merck).

GST-tagged PcyA and GST-tagged mHY2 were produced and purified as described previously (Kohchi et al., 2001; Frankenberg and Lagarias, 2003). Apo-Cph1 and apo-BphP were produced as described elsewhere (Tasler et al., 2005a; Fiege and Frankenberg-Dinkel, 2020). The concentration of the purified proteins was determined based on the absorbance at 280 nm. Extinction coefficients were calculated as described by Gill and von Hippel (Gill and von Hippel, 1989).

### Anaerobic bilin reductase assay

The anaerobic bilin reductase activity assays were performed as described previously with minor modifications (Ledermann et al., 2016). In this study, recombinant ferredoxin PetF from the cyanophage P-SSM2 was used as the electron donor in a final concentration of 1 µM. PetF was reduced using recombinant Ferredoxin-NADP^+^ reductase (FNR) from *Synechococcus* sp. PCC7002, PetH, at a final concentration of 0.01 µM (Busch et al., 2011). The reaction was started by the addition of a NADPH regenerating system consisting of 65 mM glucose-6-phosphate, 8.2 µM NADP+ and 11 U/ml glucose-6-phosphate dehydrogenase. For the assays employing defined electron equivalents, the NADPH regenerating system was replaced with a NADPH solution of a specific concentration. The photometric measurements were carried out using an Agilent 8453 spectrophotometer. The reaction was stopped by the dilution 1:10 in ice cold 0.1% TFA. Products were isolated via solid phase extraction using Sep-Pak® C18 Plus Light cartridges (Waters, Milford, Massachusetts, USA) and freeze-dried using an Alpha 2-4 LSC plus lyophilizer (Martin Christ GmbH, Osterode, Germany).

### HPLC analyses

The products of the anaerobic bilin reductase assays were analyzed using an Agilent 1100 series chromatograph equipped with a Luna® 5µm reversed phase C18-column (Phenomenex, Torrance, California, USA) and a diode-array detector. The mobile phase consisted of 50% (v/v) acetone and 50% (v/v) 20 mM formic acid at a flow rate of 0.6 mL/min. Reaction products were identified by comparison of the retention time with known standards as well as whole spectrum analysis of the elution peaks. Products of interest were collected directly at the outlet of the UV-Vis detector, frozen at -80°C and freeze-dried.

### Coupled phytochrome assembly assays

*In vitro* chromophore assembly of the apo-phytochromes Cph1 and BphP with different bilins was carried out using the lysate of a pET_*cph1* or a pASK_*bphP* overexpression after centrifugation and filtration. The assembly was tested employing 50 µL lysate incubated either with 40 µM standard bilin or with 4 µL of bilin solutions of unknown concentration for 30 min at room temperature, in the dark. Afterwards, the volume was adjusted to 500 µL with PBS. Absorbance spectra were recorded after incubation for 3 min with red light (636 nm – Pr spectrum) and after incubation for 3 min with far red light (730 nm for Cph1; 750 nm for BphP – Pfr spectrum). Difference spectra were calculated by the subtraction of the Pfr from the Pr spectrum.

### Phytofluor analyses

Phytofluor formation was evaluated via fluorescence measurements (Murphy and Lagarias, 1997). Briefly, 50 µL lysate of a pET_*cph1* or a pASK_*bphP* overexpression, after centrifugation and filtration, was incubated with 4 µL of the KflaHY2_N105D reaction product, for 30 min, at room temperature, in the dark. Afterwards, the volume was adjusted to 500 µL with PBS. Fluorescence spectra were recorded using a JASCO FP-8300 spectrofluorometer (JASCO, Tokyo, Japan).

### Protein structure prediction and analyses

KflaHY2 structure prediction was performed using AlphaFold via ColabFold (Jumper et al., 2021; Mirdita et al., 2022). Parameters set for the AlphaFold algorithm were the following: msa_mode: MMseqs2 (UniRef + Environmental), pair_mode: unpaired + paired, model_type: auto, num_recycles: 3. The obtained structure (pLDDT = 89.1) was compared to other crystalized FDBRs using the molecular visualization software PyMOL (DeLano, 2020).

### Phylogenetic analysis

A multiple sequence alignment (MSA) of ferredoxin-dependent bilin reductases was constructed using ClustalW (https://www.ebi.ac.uk/Tools/msa/clustalo/). The obtained MSA was processed employing PhyML 3.0 (http://www.atgc-montpellier.fr/phyml/) (Guindon et al., 2010) with the following command line: -D aa -B 100 -M WAG -V e -C 4 -F m -A e -O tlr. Statistical support was assessed using the transfer bootstrap expectation (TBE) in BOOSTER (https://booster.pasteur.fr/) (Lemoine et al., 2018). Tree was displayed and modified using iTOL (https://itol.embl.de/) (Letunic and Bork, 2021).

## Results

### KflaHY2 catalyzes the reduction of BV to PCB

It has been known for quite some time that the phytochromes of streptophyte algae use PCB as their chromophore instead of the PΦB found in land plants phytochrome (Wu et al., 1997; Rockwell et al., 2017). Nevertheless, the streptophyte alga *K. nitens* possesses an FDBR homolog to land plant HY2 which was indirectly suggested to be producing PCB (Rockwell et al., 2017). In order to biochemically investigate this, the reductase was recombinantly overproduced in *E. coli* and purified to test its activity (Figure 2A).

**Figure 2.**
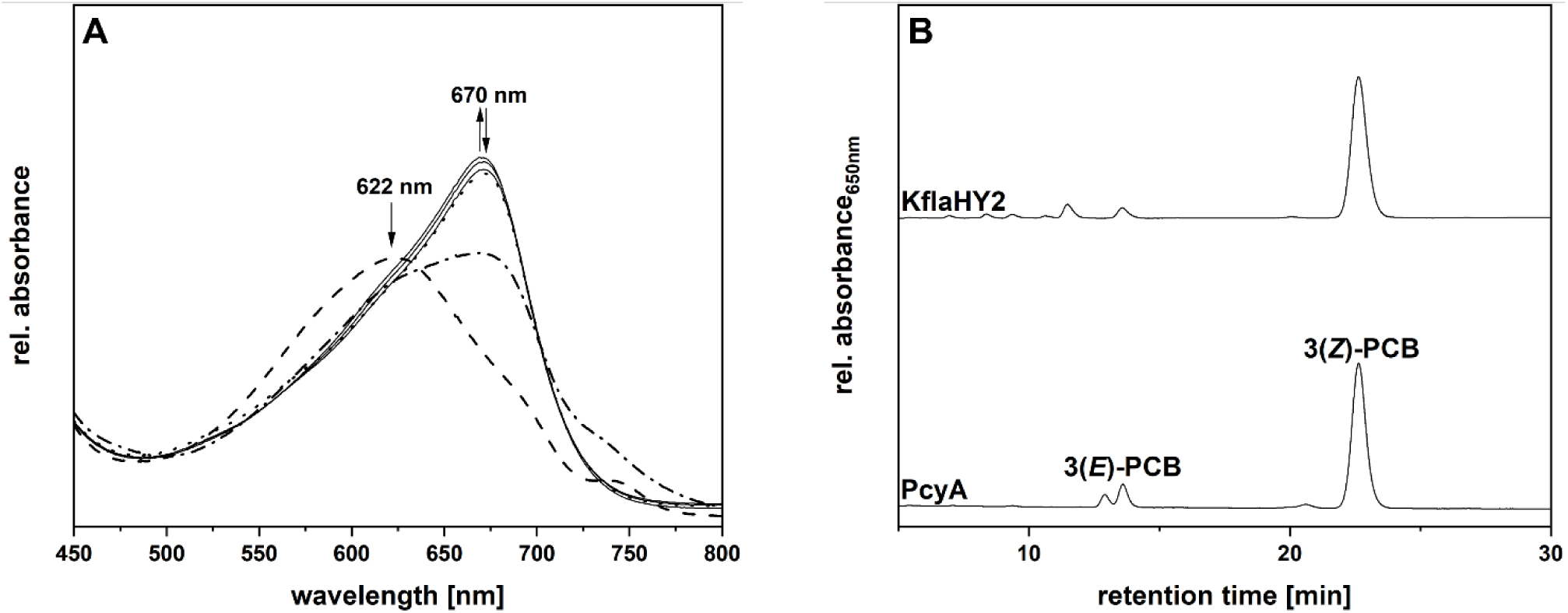
Activity assay of recombinant KflaHY2 employing BV as the substrate and characterization of reaction products. A. Time-resolved UV-Vis spectra of an anaerobic bilin reductase activity of recombinant KflaHY2 employing BV as the substrate. Total reaction time was 5 min and spectra were recorded every 30s. The course of the spectra is marked by arrows. For reasons of clarity, only relevant spectra are shown. The dashed spectrum was acquired without starting the reaction. The dotted-dashed spectrum represents the first acquired after starting the reaction. The dotted curve represents the end spectrum. B. HPLC analyses of the reaction products of KflaHY2 with BV as substrate. The products were analyzed using a reversed-phase 5 µm C18 Luna column (Phenomenex) as stationary phase. The mobile phase consisted of 50% (v/v) acetone and 50% (v/v) 20 mM formic acid flowing at 0.6 mL/min. Products of an anaerobic turnover of BV to PCB mediated by PcyA served as standards. Absorbance was continuously recorded at 650 nm.

Interestingly, the reductase and the substrate BV formed an intense turquoise colored complex with an absorbance maximum at 622 nm, suggesting that the reductase was folded correctly and was able to bind BV (Figure 2A). The presence of a shoulder at ∼ 750 nm indicates the presence of BV in its protonated form (Frankenberg and Lagarias, 2003; Tu et al., 2004). After the reaction was started, the absorbance decreased instantly. Moreover, a rapid formation of a product with an absorbance maximum at 670 nm, which can be linked to the formation of PCB, was observed. Subsequent HPLC analyses, employing the reaction products of PcyA as standards, revealed that KflaHY2 forms 3(*Z*)-PCB as the main reaction product (Figure 2B).

### KflaHY2 reaction can proceed via two intermediates

As the KflaHY2 reaction product PCB is a tetrahydro-BV, the reaction should proceed via a two-electron reduced intermediate: either the A-ring reduced PΦB, or the D-ring reduced 18^1^,18^2^-DHBV. Intermediates of the reaction were isolated using the products of an anaerobic bilin reductase activity assay conducted with one electron equivalent of NADPH. HPLC analyses of the products showed that KflaHY2 forms two intermediates (Figure 3 – marked as “Inter 1” and “Inter 2”).

**Figure 3.**
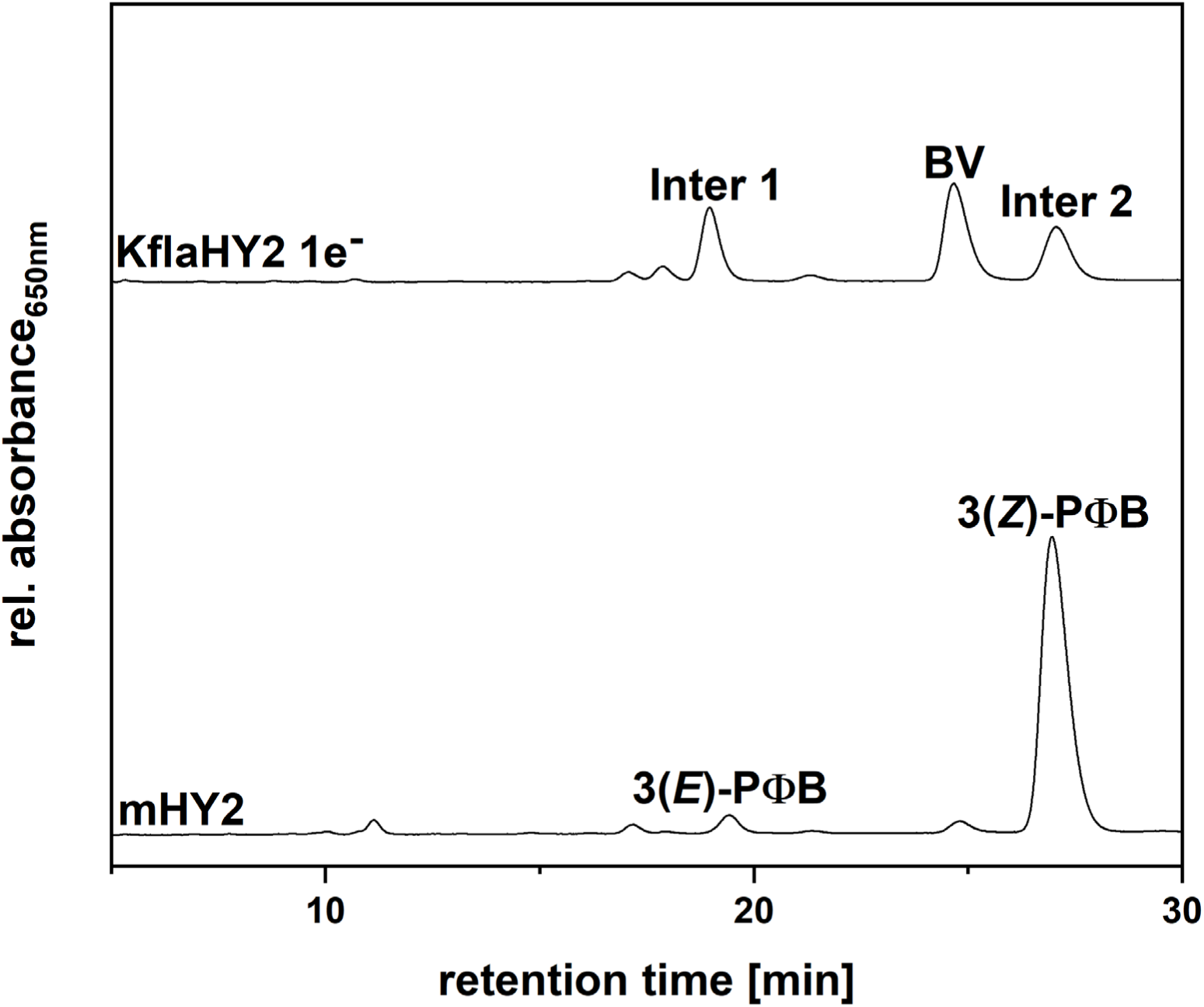
HPLC analyses of the reaction products of an anaerobic activity assay employing equimolar amounts of KflaHY2 and BV with one electron equivalent of NADPH. Products were resolved using a reversed-phase 5 µm C18 Luna column (Phenomenex) as stationary phase and a mixture of 50% (v/v) acetone and 50% (v/v) 20 mM formic acid as mobile phase at 0.6 mL/min. The products of an anaerobic turnover of BV to PΦB mediated by HY2 derived from *A. thaliana* were used as standards. Absorbance was continuously recorded at 650 nm.

The intermediates possessed the same retention times as the products 3(*E*)-PΦB (∼ 19 min) and 3(*Z*)-PΦB (∼ 27 min) of the recombinant reductase HY2 from *A. thaliana* (mHY2). However, 18^1^,18^2^-DHBV, the intermediate of the cyanobacterial PcyA reaction towards PCB, and 3(*E*)-PΦB are roughly characterized by the same retention time. Therefore, “Inter 1” could have been either of the two (Frankenberg and Lagarias, 2003).

### The intermediates formed by KflaHY2 are 18^1^,18^2^-DHBV and 3(*Z*)-PΦB

To confirm the nature of the intermediates identified by HPLC analyses, coupled phytochrome assembly assays of the isolated intermediates with the apo-phytochromes Cph1 from *Synechocystis* sp. PCC 6803 and BphP from *Pseudomonas aeruginosa* were conducted (Figure 4A). Cph1 finds its natural chromophore in PCB, while BphP uses BV. However, phytochromes are also able to accommodate different bilins as long as they possess the same A-ring configuration as the natural chromophore. Thus, Cph1 is able to bind bilins possessing an ethylidene group at the A-ring, whereas BphP only binds bilins with an A-ring endo-vinyl group (Li and Lagarias, 1992; Tasler et al., 2005b). Apo-Cph1 incubated with “Inter 1” did not produce a photoactive adduct since no characteristic difference spectrum was obtained, indicating the lack of the ethylidene group at the A-ring C3 side chain (Figure 4A – dashed line) (Xu et al., 2019). This led to the conclusion that “Inter 1” could not be 3(*E*)-PΦB. Incubation of apo-Cph1 with “Inter 2” (Figure 4A – dotted line) led to a difference spectrum like it was obtained for the incubation of Cph1 with 3(*Z*)-PΦB (Figure 4A – solid line). These results, as well as the results of the HPLC analyses (Figure 3), prove that “Inter 2” is 3(*Z*)-PΦB.

**Figure 4.**
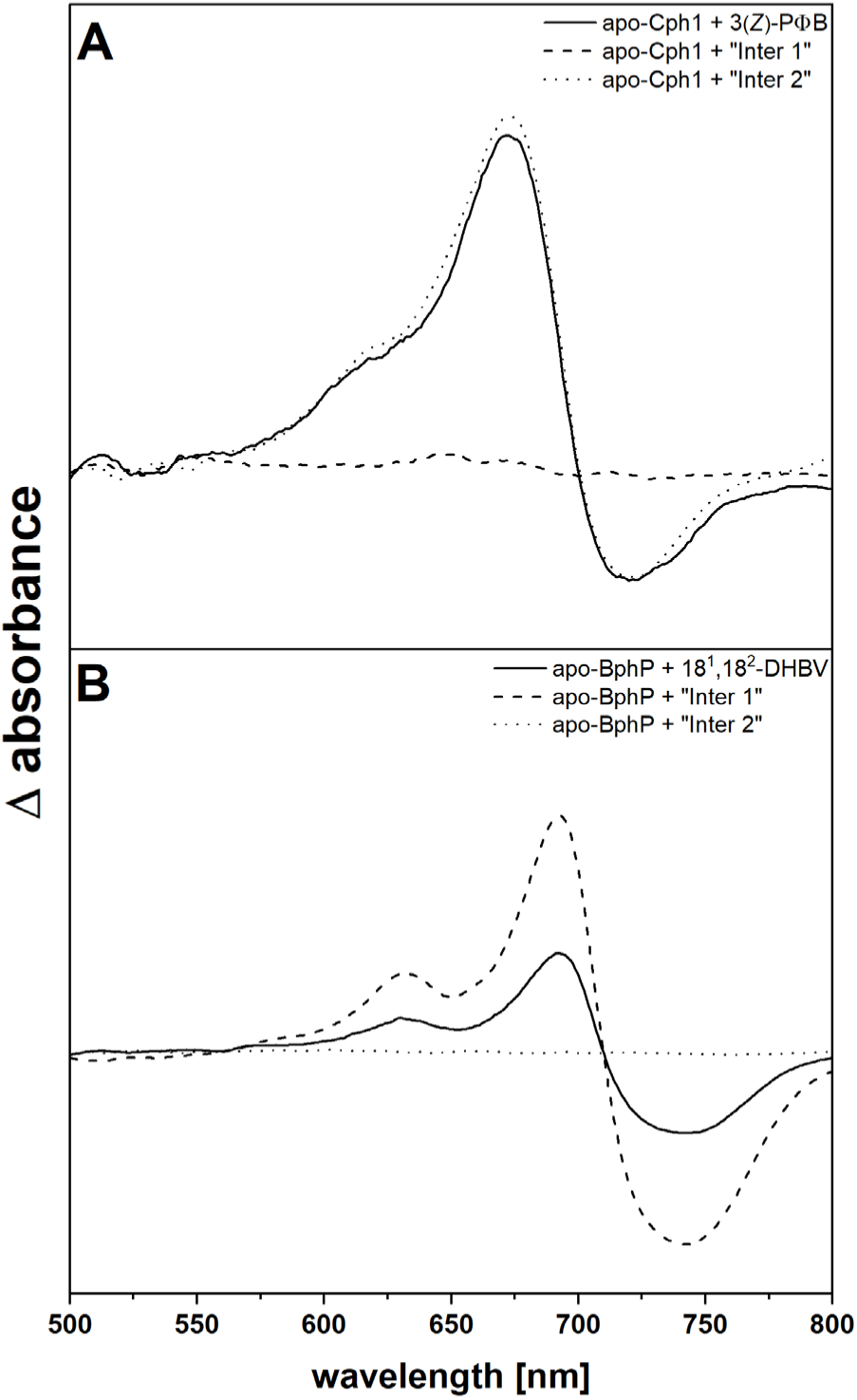
Red/far-red light induced difference spectra of coupled phytochrome assembly assays employing the intermediates of the KflaHY2 reaction. Absorbance spectra were recorded after incubation for 3 min with red light (636 nm – Pfr spectrum) and after incubation for 3 min with far red light (730 nm for Cph1; 750 nm for BphP – Pr spectrum). Difference spectra were calculated by the subtraction of the Pfr from the Pr spectrum. All calculated difference spectra were smoothed applying a 20 pt. Savitzky-Golay filter. A. Coupled phytochrome assembly assays employing apo-Cph1 and “Inter 1” (dashed line); apo-Cph1 and “Inter 2” (dotted line); apo-Cph1 and 3(*Z*)-PΦB (solid line) B. Coupled phytochrome assembly assay employing apo-BphP and “Inter 1” (dashed line); apo-BphP and “Inter 2” (dotted line); apo-BphP and 18^1^,18^2^-DHBV (solid line).

On the other hand, “Inter 2” failed to form a photoactive adduct with apo-BphP (Figure 4B – dotted line), whereas, “Inter 1” formed a photoactive adduct with apo-BphP with a difference spectrum (Figure 4B – dashed line) similar to the difference spectrum obtained by the incubation of BphP with 18^1^,18^2^-DHBV (Figure 4B – solid line). This revealed that “Inter 1” is indeed 18^1^,18^2^-DHBV.

To clarify whether the intermediates are only artifacts of the *in vitro* assay or if they are actually suitable in KflaHY2-catalyzed BV reduction, both compounds were used as substrates for KflaHY2 in anaerobic bilin reductase activity assays. Subsequent HPLC analyses revealed that both intermediates are transformed to mostly 3(*Z*)-PCB (Figure 5).

**Figure 5.**
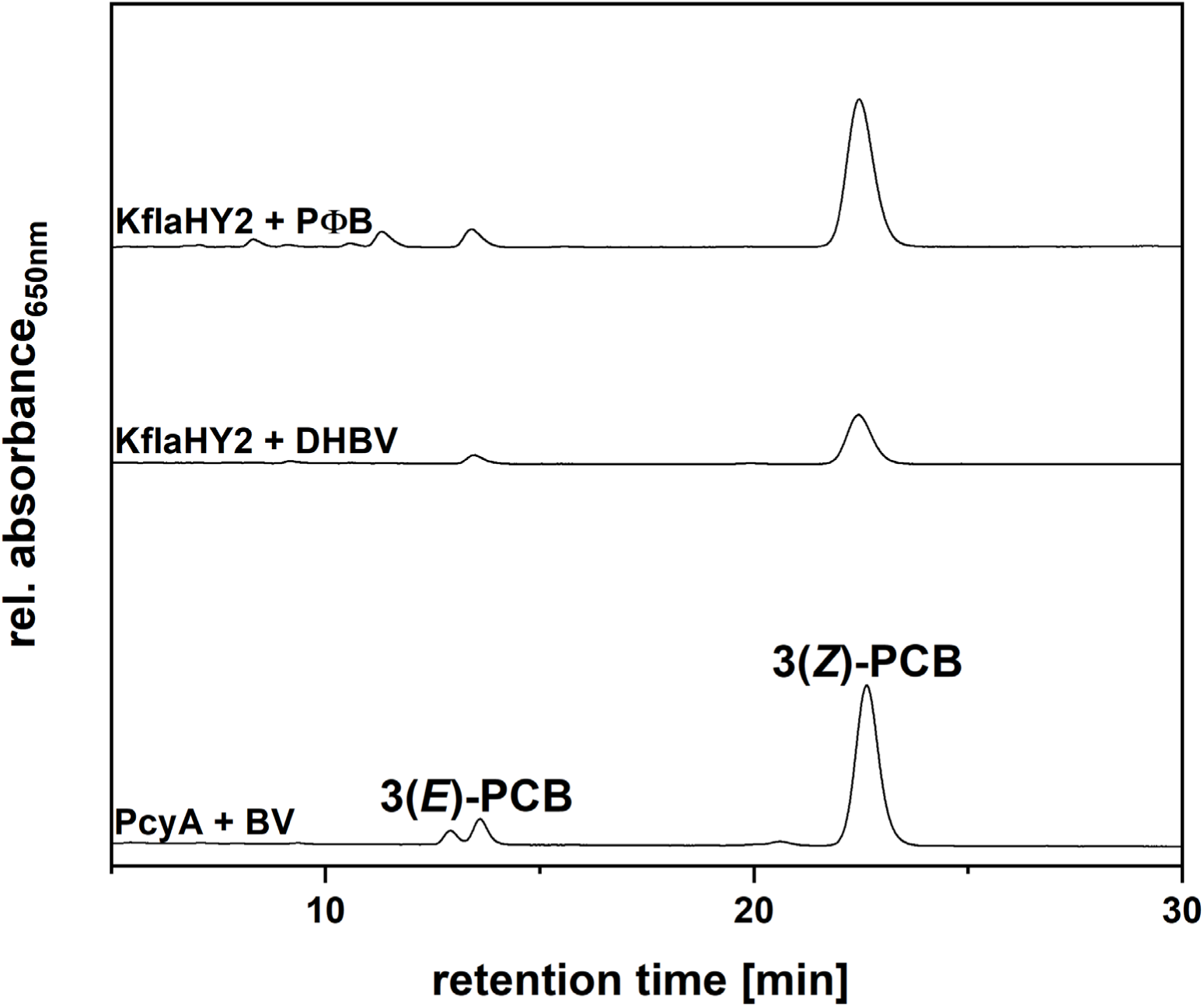
HPLC analyses of the reaction products of activity tests employing 3(*Z*)-PΦB and 18^1^,18^2^-DHBV as substrates for KflaHY2. Products were analyzed using a 5 µm reversed-phase C18 Luna column (Phenomenex) as stationary phase. The mobile phase consisted of 50% (v/v) acetone and 50% (v/v) 20 mM formic acid flowing at 0.6 mL/min. Reaction products of a PcyA mediated reduction of BV to PCB were used as standards. Absorbance was continuously measured at 650 nm.

### Asp^122^ and Asp^242^ are essential for the catalytic activity

In order to gain further insight on the unexpected behavior of KflaHY2 and to learn about the evolution of HY2, an amino acid sequence alignment employing different FDBRs was constructed (Figure S2). Three residues caught our attention: Asn^105^, Asp^122^ and Asp^242^. KflaHY2 Asn^105^ represents the homolog to *Synechocystis sp.* PCC 6803 PcyA His^88^, which was proven to be involved in protonation and substrate positioning in the active site (Hagiwara et al., 2010). The substitution of the His to an Asn is not unusual since it can also be found in other FDBRs like PebA, PebB and PebS. On the other hand, land plants HY2s show an Asp in the corresponding position which was also proven to be essential for substrate positioning in *Arabidopsis thaliana* HY2 and involved in catalysis in *Solanum lycopersicum* HY2 (SlHY2) (Tu et al., 2008; Sugishima et al., 2020). KflaHY2 Asp^122^ finds a homolog in all the FDBRs except for the ones from land plants. In PcyA and PebS, it is located in the central β-sheet of the binding pocket and was shown to be the initial proton donor (Hagiwara et al., 2006; Busch et al., 2011). Finally, Asp^242^ finds a homolog in Asp^256^ of *A. thaliana* HY2, Asp^256^ of *S. lycopersicum* HY2 (SlHY2), Asp^206^ of PebS and Asp^219^ of *Guillardia theta* PEBB. All of these Asp residues were shown to be involved in the reduction of the A-ring 2,3,3^1^,3^2^-diene system (Tu et al., 2008; Busch et al., 2011; Sommerkamp et al., 2019; Sugishima et al., 2020). On the basis of these considerations, site-directed mutagenesis was performed. To investigate the influence of ionization on catalysis, Asp^122^ and Asp^242^ were replaced by asparagine residues whereas Asn^105^ was replaced by an aspartate residue. Moreover, two double variants were generated. In particular, one was constructed for KflaHY2 to resemble a plant HY2, the other to prove the presence of two Asp residues is enough for reducing BV to PCB. The activity of the purified variant enzymes was investigated *in vitro* via FDBR assays and subsequent HPLC analyses (Figure 6).

**Figure 6.**
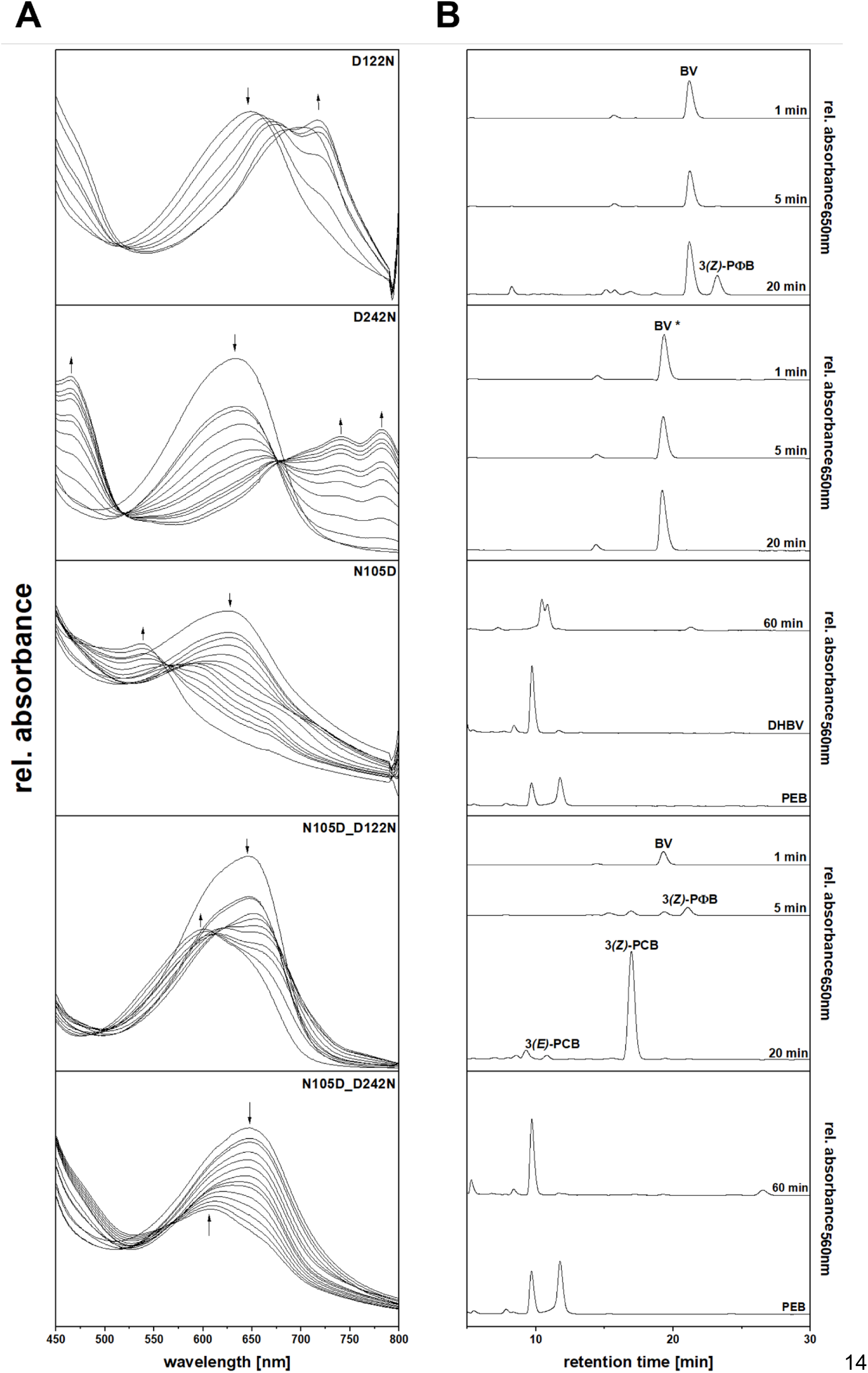
Activity assay and characterization of reaction products of recombinant KflaHY2 variants. A. UV-Vis spectra of the anaerobic FDBR Assay performed using KflaHY2 variants and BV as the substrate. Total reaction time was 20 minutes for KflaHY2_D122N, KflaHY2_D242N, KflaHY2_N105D_D122N and 60 min for KflaHY2_N105D, KflaHY2_N105D_D242N. Spectra were recorded every 30 seconds. For reasons of clarity, only relevant spectra are shown. The arrow indicates the course of absorbance during the reaction. B. HPLC elution profiles of the reaction products of KflaHY2 variants. The stationary phase consisted in a Luna 5µm reversed phase C18 column (Phenomenex). The mobile phase consisted of a mixture of 50% (v/v) acetone and 50% (v/v) 20mM formic acid flowing at 0.6 mL/min. Absorbance was continuously recorded at 560 or 650 nm. BV shows change in the retention time due to substitution of the mobile phase between the different HPLC measurements (*).

The D122N variant was able to bind BV and use it as a substrate, since the formation of a peak at ∼ 720 nm was observed (Figure 6A – D122N). Intermediates were isolated from the reaction mix at certain time points and analyzed via HPLC. Time-course HPLC analyses revealed the formation of small amounts of 3(*Z*)-PΦB after 20 min from the start of the reaction (Figure 6B – D122N). This variant partially retains the capability to reduce the BV A-ring but is completely unable to reduce the D-ring, thus proving the importance of D122 for D-ring reduction.

The D242N variant still retained the ability to bind BV but failed in its reduction. The increase of the absorption at ∼ 460, ∼ 740 and ∼ 790 nm suggested the formation of the radical BVH• (Figure 6A – D242N) (Tu et al., 2004; Tu et al., 2008; Busch et al., 2011). The HPLC elution profiles were compatible with BV, therefore confirming the inactivity (Figure 6B – D242N). This result is consistent with what was found for mHY2_D256N and SlHY2_D263N indicating the importance of this residue for BV protonation (Tu et al., 2008; Sugishima et al., 2020).

The N105D variant showed an unexpected behavior. The assay was initially performed for 20 min but strangely revealed the formation of a violet-colored product with a peak ∼ 570 nm, suggesting production of either phycoerythrobilin (PEB) or 15,16-dihydrobiliverdin (15,16-DHBV) (data not shown). In order to potentially accumulate more of this violet compound, we decided to let the reaction run for a longer time, 60 min. This second assay showed the formation of a final product with an absorbance peak at ∼ 540 nm (Figure 6A – N105D). Time-based HPLC analyses revealed that the bilin produced after 5 min from the start of the reaction was indeed PCB, whereas a pattern compatible with the unidentified compound and mostly 3(*Z*)-PCB was obtained for the 20 min product (data not shown). As for the final 60 min product, HPLC analysis revealed a double peak with the retention time of 10.5 and 10.8 min, not matching the retention time of any known bilin and therefore leading to uncertainty (Figure 6B – N105D). To identify the unknown bilin, its integration in the phytochromes Cph1 and BphP was evaluated. PEB and 15,16-DHBV lack the C15-16 double bond required for the photoconversion of the phytochrome between the two isoforms. Therefore, phytochromes binding bilins lacking the C15-16 double bond are locked in the red-light absorbing form. In this condition, there is no complete de-excitation of the bilin, leading to the formation of a fluorescent product named phytofluor (Murphy and Lagarias, 1997). The bilin product isolated from the reaction catalyzed by KflaHY2_N105D was then incubated in parallel with the two apo-phytochromes and fluorescence was measured. The formation of a characteristic phytofluor was only observed for the incubation with apo-Cph1, thus suggesting the product is PEB (Figure 7).

**Figure 7.**
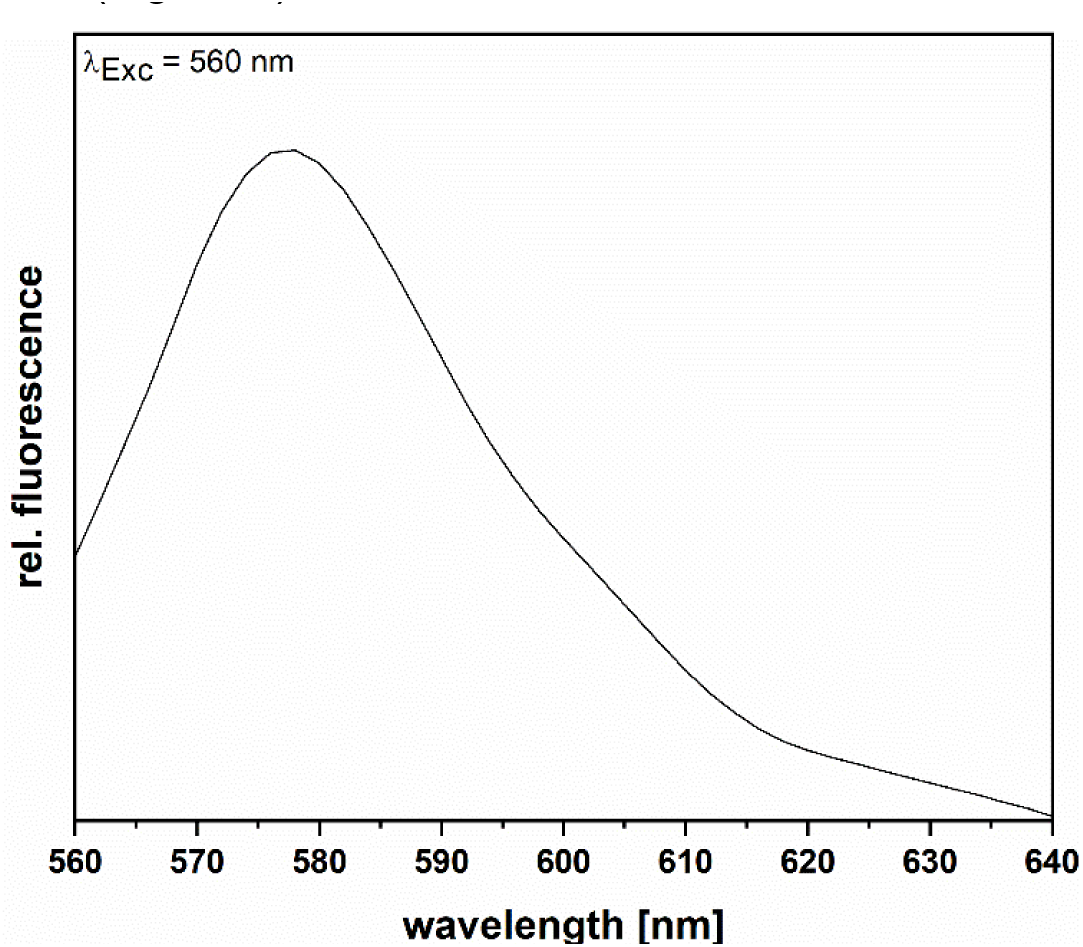
Fluorescence emission spectrum of the apo-Cph1-KflaHY2_N105D reaction product adduct. The isolated bilin produced by KflaHY2_N105D was incubated for 30 min in the dark with the lysate of a pET_*cph1* overproduction. Fluorescence emission was recorded after excitation at 560 nm.

The double variant N105D_D122N, constructed to mimic a plant HY2, surprisingly retained the native activity, revealing the conversion of BV to PCB instead of the hypothesized 2e^-^ reduction to PΦB (Figure 7 – N105D_D122N). This result led to the hypothesis that the presence of two aspartate residues in the active site, even if in different position than in the native enzyme, is enough for the reduction of BV to PCB. To further investigate it, the double variant N105D_D242N was also generated. Interestingly, performing the assay for 60 min revealed the formation of a pink product with an absorbance at ∼ 600 nm (Figure 6A – N105D_D242N). Via HPLC analysis, the final product was identified to be 3(*E*)-PEB (Figure 6B – N105D_D242N), thus showing a different activity than expected.

### *Netrium digitus* HY2 is also a PCB synthesizing enzyme

Since our attempt to convert the *K. nitens* HY2 enzyme into a plant type HY2 with PΦB biosynthetic activity failed, we decided to go back to a phylogenetic tree of the HY2 lineage (Figure 8) to select sequences for biochemical assays and identify where the change of activity happened.

**Figure 8.**
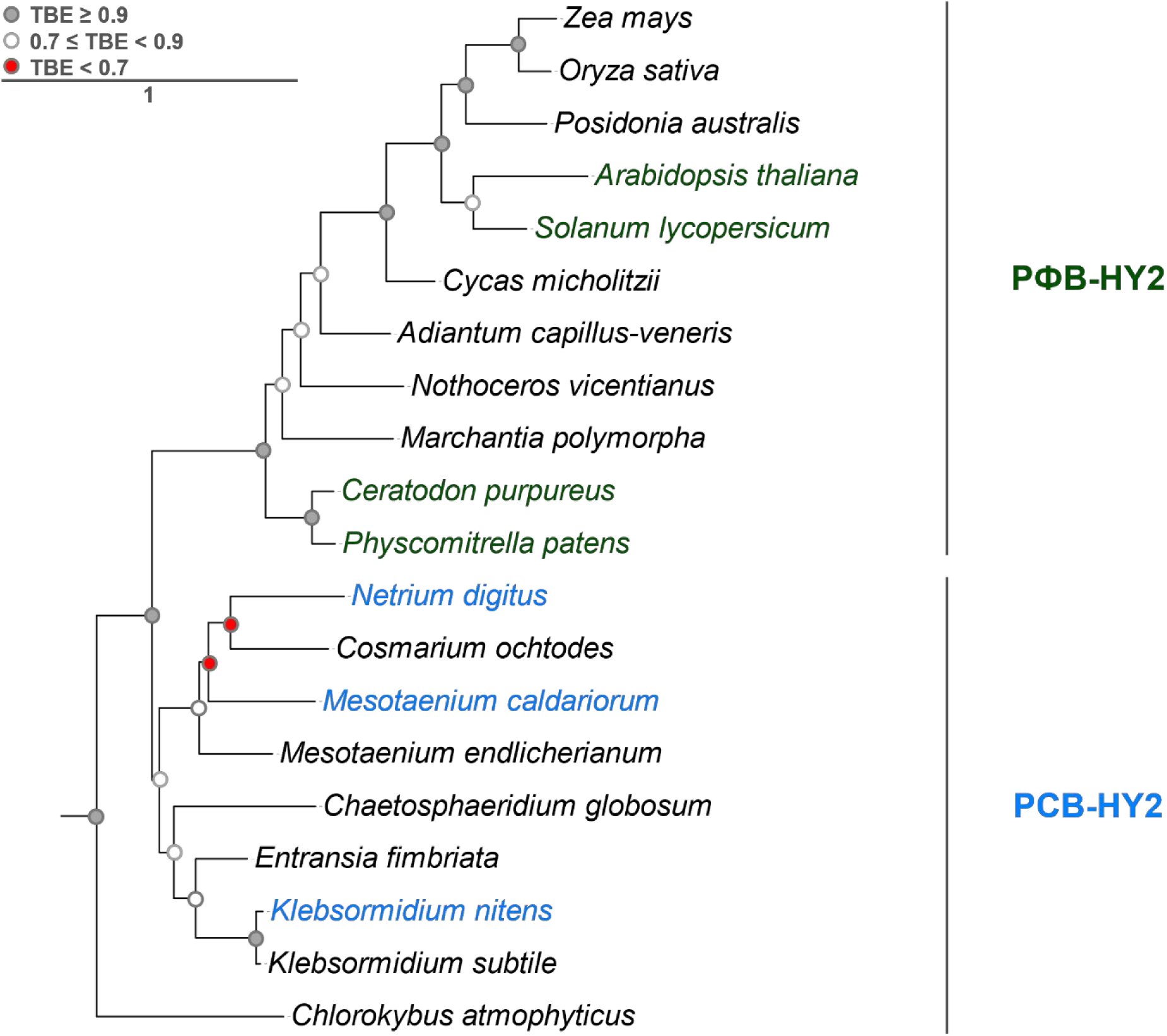
Phylogenetic analysis of HY2 lineage. The maximum-likelihood tree was inferred using PhyML (Guindon et al., 2010). Transfer bootstrap expectation (TBE) was used for support. The represented tree is an excerpt of a full tree including members of all the FDBRs lineages (Figure S). Accession numbers of the employed sequences shown in this figure are available in Supplement material (Table S3). Labels indicate HY2-belonging organism. The color of the label represents the produced bilin (cyan for PCB, dark green for PΦB). Only HY2 with experimentally proven activity are colored. Circles at nodes indicate TBE (grey, TBE ≥ 0.9; white, 0.7 ≤ TBE < 0.9; red, TBE < 0.7).

The *Physcomitrella patens* HY2 has been studied previously and was shown to convert BV to PΦB (Chen et al., 2012) (Shih-Long Tu, Academia Sinica, Taiwan, personal communication). *Mesotaenium caldariorum* was previously shown to synthesize its PCB phytochrome chromophore via PФB using cell free extracts (Wu et al., 1997). Therefore, we decided to have a look at the HY2s from *Ceratodon purpureus* and *Netrium digitus* (NediHY2). Unfortunately, both recombinant proteins were prone to aggregation but we managed to obtain enough catalytic activity for product identification. While the HY2 of *C. purpureus* confirmed previous indirect evidence and synthesized PΦB from BV (Lamparter et al., 1995) (data not shown), NediHY2 was shown to produce PCB in a coupled phytochrome assay using apo-Cph1 (Figure 9).

**Figure 9.**
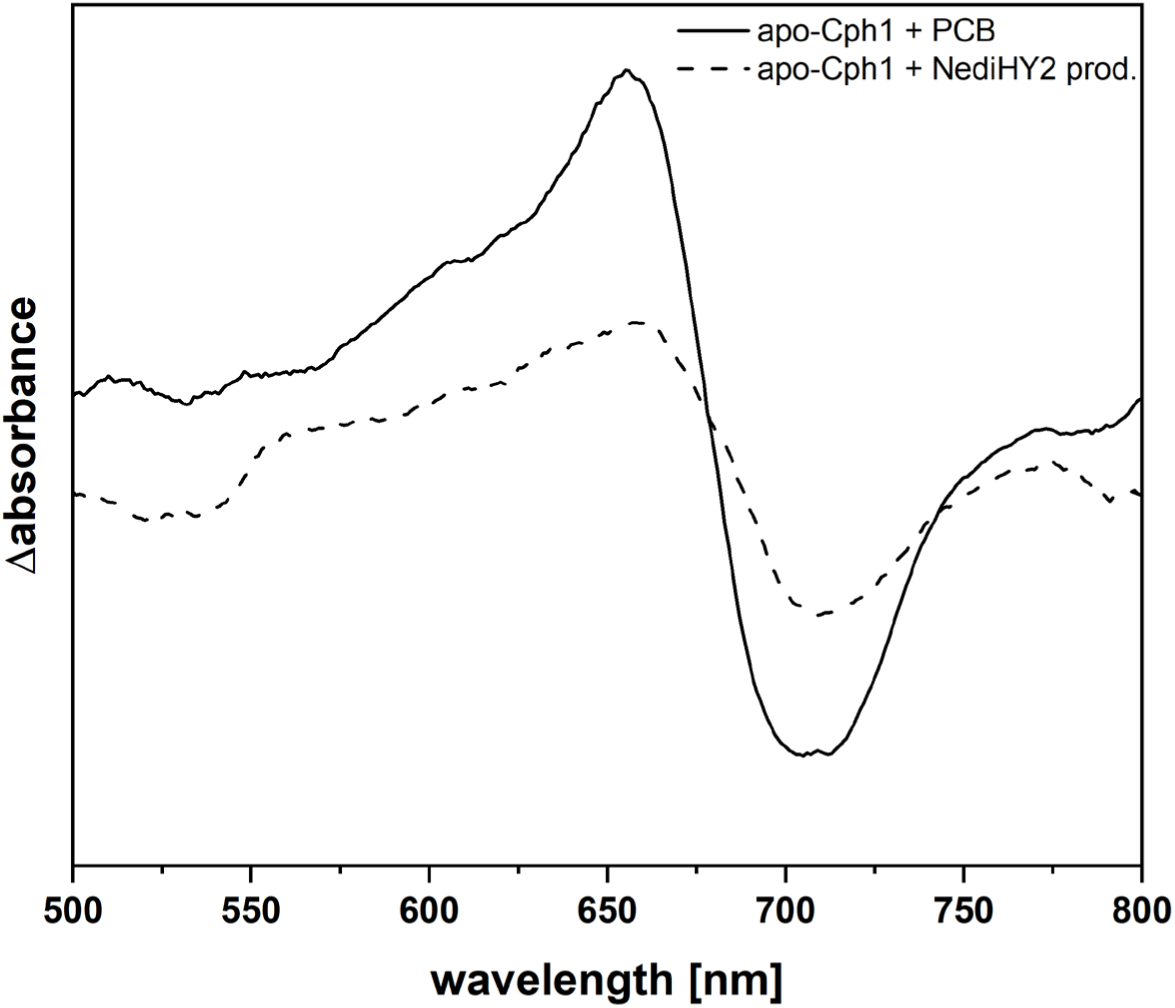
Red/far-red light induced difference spectra of coupled phytochrome assembly assays employing PCB and the product of the NediHY2 reaction. Absorbance spectra were recorded after incubation for 3 min with red light (636 nm – Pfr spectrum) and after incubation for 3 min with far red light (730 nm – Pr spectrum). Difference spectra were calculated by the subtraction of the Pfr from the Pr spectrum. The apo-Cph1 + PCB difference spectrum (solid line) was smoothed applying a 20 pt. Savitzy-Golay filter. The apo-Cph1 + NediHY2 prod. difference spectrum (dashed line) was smoothed applying a 80 pt. Savitzky-Golay filter.

Based on these data and the phylogenetic tree of the HY2 lineage of FDBRs, we can conclude that the switch of activity from a PCB-producing to a PΦB-producing HY2 enzyme happened within the streptophyte HY2 lineage, likely between *Netrium digitus* and *Physcomitrella patens*.

## Discussion

Over two decades ago, the last of the classic photomorphogenetic mutants of *Arabidopsis thaliana*, *hy2*, was cloned (Kohchi et al., 2001). *HY2* encodes phytochromobilin synthase, the first member of the ferredoxin-dependent bilin reductase family, catalyzing the last step in plant phytochrome chromophore biosynthesis. Subsequently, multiple members of the FDBR family were identified in cyanobacteria, red algae, mosses and cryptophytes (Frankenberg et al., 2001; Chen et al., 2012; Overkamp et al., 2014; Rockwell and Lagarias, 2017). All share the same overall fold consisting of an α-/β-/α-sandwich which harbours the active side on top of the central β-sheet. With the number of recombinant FDBRs biochemically characterized, it became clear that it is not always possible to predict the catalytic activity solely based on an amino acid sequence alignment (Dammeyer et al., 2008; Ledermann et al., 2018). One reason is attributable to the substrate binding mode. Indeed, the crystal structure of both PebB and HY2 revealed a *ZZZssa* flipped binding mode of the linear tetrapyrrole substrate (Sommerkamp et al., 2019; Sugishima et al., 2020). Both enzymes have furthermore in common that they reduce the A-ring 2,3,3^1^,3^2^-diene system of the substrate. Their relatedness is phylogenetically supported as HY2 is part of the PebB lineage of FDBRs (Rockwell and Lagarias, 2017). That said, it was rather unexpected to find out that streptophyte algae all bare HY2 homologs, although their phytochromes employ a PCB chromophore (Wu et al., 1997; McDowell and Lagarias, 2001; Rockwell et al., 2017). These bilins are characterized by different spectral properties: while PCB absorbs orange and red light (λ_PCB_ = ∼ 620 - 670 nm), PΦB absorbs far-red light (λ_PΦB_ = ∼ 660 - 685 nm). In the present study, we confirmed that the HY2 of *Klebsormidium nitens* is indeed a functional FDBR, which, in contrast to the phylogenetic classification as a HY2, possesses a PcyA-like activity, catalyzing the reduction of BV to PCB. Furthermore, we identified 18^1^,18^2^-DHBV and 3(*Z*)-PΦB as intermediates occurring under the chosen assay conditions. These results were somehow unexpected, as they lead to the conclusion that KflaHY2 can mediate the reduction of BV to PCB employing two different reaction routes (Figure 1). However, the formation of 3(*Z*)-PΦB as an intermediate during PCB biosynthesis is tying well with previous phytochrome studies in the Zygnematophycea *Mesotaenium caldariorum*, an organism that also possesses a HY2 (Figure 9) (Wu et al., 1997). Taking this into consideration, we propose that PΦB is the real intermediate in the KflaHY2 catalyzed conversion of BV to PCB and the formation of the 18^1^,18^2^-DHBV intermediate is rather an experimental artefact. Here it is possible that, during *in vitro* enzyme assays, the substrate BV is forced into the active site in two orientations leading to the two observed isomers. *In vivo*, the substrate might be directly channeled from the preceding enzyme, heme oxygenase. A similar scenario has been described for certain heme oxygenases from pathogenic bacteria where the orientation of the substrate dictates the product formed (Caignan et al., 2002). This is further supported by a structural model of KflaHY2 obtained using AlphaFold (Jumper et al., 2021) and two structural alignments using PyMOL: one with the crystal structure of the HY2 of *Solanum lycopersicum*, with which KflaHY2 shares high sequence homology, the second with the crystal structure of PcyA of *Synechocystis* sp. PCC 6803 (SynPcyA), catalyzing BV to PCB reduction as KflaHY2 (Figure 10) (Hagiwara et al., 2006; DeLano, 2020; Sugishima et al., 2020). The obtained alignments revealed a higher similarity of the KflaHY2 active site to the one of SlHY2. Critical residues of the bottom β-sheet superimpose perfectly, and also the distal Asp residue (SlHY2-Asp^263^/KflaHY2-Asp^242^), important for A-ring reduction, is placed similarly (Figure 10A). In contrast, the corresponding residue in SynPcyA (Asp^220^) is facing outside of the active site and was proven to be not involved in catalysis (Figure 10B) (Hagiwara et al., 2006). On the other hand, the reduction of the D-ring exo-vinyl group in SynPcyA was proven to be catalyzed by the residue Glu^76^, which does not find a homolog in KflaHY2 (Hagiwara et al., 2006). Based on our biochemical data, Asp^122^ in KflaHY2 could fulfill this function.

**Figure 10.**
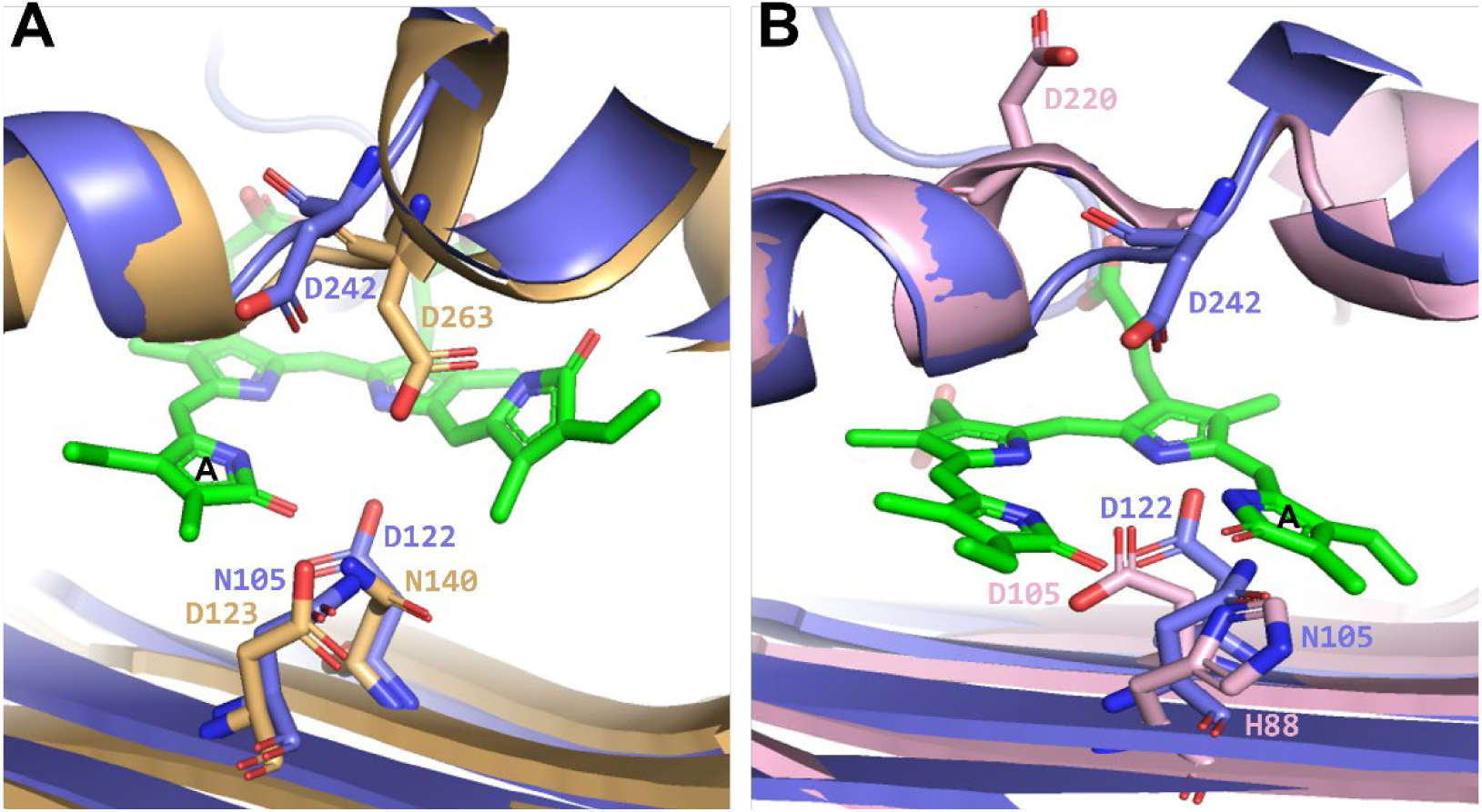
Structures alignment of KflaHY2 with SlHY2 and SynPcyA. The model structure of KflaHY2 (displayed in slate cartoon) was obtained using AlphaFold. The alignments were generated using PyMOL (DeLano, 2020). A. Overlay of the active sites of KflaHY2 and SlHY2. The crystal structure of SlHY2 (PDB accession code: 6KME) is displayed as gold cartoon. Biliverdin IXα, in the conformation assumed in the active site of SlHY2, is shown in green, with A-ring being labeled. Amino acid residues of interest are shown as labeled sticks and colored as the belonging structure. B. Overlay of the active sites of KflaHY2 and SynPcyA. The crystal structure of SynPcyA (PDB accession code: 2D1E) is displayed as pink cartoon. Biliverdin IXα, in the conformation assumed in the active site of SlHY2, is shown in green, with A-ring being labeled. Amino acid residues of interest are shown as labeled sticks and colored as the belonging structure.

The specificity of reduction is furthermore influenced by the substrate binding mode of BV, shown to be flipped in the PebB/HY2 lineage of FDBRs (Sommerkamp et al., 2019; Sugishima et al., 2020). We propose this is also true for KflaHY2 as two critical amino acid residues responsible for the stabilization of BV in the *ZZZssa* flipped binding mode are conserved. Overall, the data indicate that the ability of KflaHY2 to synthesize PCB has evolved independently from the biosynthesis catalyzed by PcyA.

Surprisingly, our attempt to convert KflaHY2 into a land plant HY2 by exchanging the catalytic pair was unsuccessful in yielding PΦB. We rather revealed that an additional proton donating residue on the β-sheet of the active site induced isomerization of the product to PEB (Figure S3). Product specificity in the HY2 lineage therefore could rather be determined by the architecture of the active site and other residues in the surroundings of the binding pocket. This became even clearer when studying the FDBRs from *Netrium digitus* and *Ceratodon purpureus*. Both FDBRs share with KflaHY2 about 48% sequence identity and the same catalytic pair. While the HY2s of *N. digitus* and *K. nitens* are PCB-HY2s and group well with the other characterized HY2 from *Mesotaenium caldariorum* in the phylogenetic tree, the *C. purpureus* enzyme is a PΦB-HY2 and forms a monophyletic group with the *Physcomitrella patens* HY2 and land plant HY2s (Figure 9). Based on this, we propose that within the HY2 lineage, two monophyletic groups diverged from the common HY2 ancestor: the PCB-HY2 clade and the PΦB-HY2 clade. Putting these data into perspective, one would therefore suggest that phytochromes with a PΦB chromophore were first found in the bryophytes and that early diverging streptophyte like *Klebsormidium* still possessed a PCB-containing phytochrome. This might be related to the available wavelength of light to these organisms, as a PCB-containing phytochrome will absorb more in the red/orange region of the visible spectrum compared to that a PΦB-containing phytochrome. While these phytochromes are thought to have a common cyanobacterial ancestor, the corresponding FDBR sequences have significantly diversified since primary endosymbiosis (Rockwell and Lagarias, 2017).

## Acknowledgement

This project was financially supported by a grant from the Deutsche Forschungsgemeinschaft to NFD and EH. FZ and EH acknowledge support by the Deutsche Forschungsgemeinschaft within the Research Training Group GRK2341 “MiCon”. We furthermore thank J. Clark Lagarias for the gift of plasmids and helpful discussions with him and Nathan Rockwell.

## Author contributions

FF, BL and NFD designed research, FF, BL, JH, ZF performed research, EPP and EH contributed analytical tools, FF, BL and NFD wrote the paper. All authors read and approved the final version of the manuscript.

