## supplemental material for "On the evolution of the plant phytochrome chromophore biosynthesis"

**Table S1** List of employed primers for Quikchange mutagenesis. Only forward primers are given, reverse primers are the complement

| Primer | Sequence (5'→3') |
|---|---|
| **pET28a_*KflaHY2*_N105D_fwd** | CAATCTGCAAGTTCTGGATCTGATCGCGTTTCCG |
| **pET28a_*KflaHY2*_D122N_fwd** | CGTACTTCTGTGCAAACCTGGTCACCCTG |
| **pET28a_*KflaHY2*_D242N_fwd** | GGCGTGCAGAAAAAAATCCGGGTCGTCCG |

**Table S2** List of employed constructs

| Plasmid | Features | Accession number | Reference |
|---|---|---|---|
| **pET28a_*KflaHY2*** | pET-28a(+) derivate carrying a synthetic gene comprising the catalytic core region of KflaHY2 lacking the predicted chloroplast transit peptide; N-terminal $His_6$-tag | A0A1Y1I5X3 (UniProt) | Donation of J. Clark Lagarias, UC Davies (unpublished) |
| **pET28a_*KflaHY2*_N105D** | pET28a_KflaHY2 derivate obtained by site-directed mutagenesis | | This study |
| **pET28a_*KflaHY2*_D122N** | pET28a_KflaHY2 derivate obtained by site-directed mutagenesis | | This study |
| **pET28a_*KflaHY2*_D242N** | pET28a_KflaHY2 derivate obtained by site-directed mutagenesis | | This study |
| **pET28a_*KflaHY2*_N105D_D122N** | pET28a_KflaHY2 derivate obtained by site-directed mutagenesis | | This study |
| **pET28a_*KflaHY2*_N105D_D242N** | pET28a_KflaHY2 derivate obtained by site-directed mutagenesis | | This study |
| **pGEX_*pcyA*** | pGEX-6P-1 derivate carrying the *pcyA* gene from *Nostoc* sp. PCC 7120; N-terminal GST-tag | Q93TN0 (UniProt) | Frankenberg et al., 2001 |

| | | | |
|---|---|---|---|
| **pGEX_*mHY2*** | pGEX-6P-1 derivate carrying the mature *HY2* gene from *Arabidopsis thaliana*; N-terminal GST-tag | Q9SR43 (UniProt) | Kohchi et al., 2001 |
| **pGEX_*petF*_P-SSM2** | pGEX-6P-3 derivate carrying a synthetic gene of the *petF* gene from the cyanophage P-SSM2; N-terminal GST-tag | Q58M74 (UniProt) | Dammeyer et al., 2008 |
| **pGEX_*petH*** | pGEX-6P-1 derivate carrying the *petH* gene from *Synechococcus* sp. PCC 7002; N-terminal GST-tag | P31973 (UniProt) | Unpublished, lab collection |
| **pET_*cph1*** | pET derivate carrying the *cph1* gene from *Synechocystis* sp. PCC6803 | Q55168 (UniProt) | Donation of J. Clark Lagarias, UC Davis |
| **pASK_*bphP*** | pASK-IBA3 derivate carrying the *bphP* gene from *Pseudomonas aeruginosa* | Q9HWR3 (UniProt) | Tasler *et al*, 2005 |
| **pGEX-4T-1_*CepuHY2*** | pGEX-4T-1 derivate carrying the HY2 gene from C*eratodon purpureus* | FFPD-2057880 (OneKP) | This study |
| **pGEX-4T-1_*NediHY2*** | pGEX-4T-1 derivate carrying the HY2 gene from *Netrium digitus* | FFGR-2000050 (OneKP) | This study |

**Table S3** List of HY2 sequences used for the construction of the phylogenetic tree

| **Organism** | **Database** | **Accession number** |
|---|---|---|
| *Chlorokybus atmophyticus* | OneKP | AZZW-2019954 |
| *Klebsormidium subtile* | OneKP | FQLP-2031011 |
| *Klebsormidium nitens* | GenBank | GAQ84116.1 |
| *Entransia fimbriata* | OneKP | BFIK-2028189 |
| *Chaetosphaeridium globosum* | OneKP | DRGY-2006866 |
| *Mesotaenium endlicherianum* | OneKP | WDCW-2047152 |
| *Mesotaenium caldariorum* | OneKP | HKZW-2005125 |
| *Cosmarium ochtodes* | OneKP | STKJ-2007533 |
| *Netrium digitus* | OneKP | FFGR-2000050 |
| *Physcomitrella patens* | GenBank | BAF02520.1 |
| *Ceratodon purpureus* | OneKP | FFPD-2057880 |
| *Marchantia polymorpha* | OneKP | JPYU-2037290 |

| | | |
|---|---|---|
| *Nothoceros vicentianus* | OneKP | TCBC-2082726 |
| *Adiantum capillus-veneris* | GenBank | BAF02518.1 |
| *Cycas micholitzii* | OneKP | XZUY-2012311 |
| *Solanum lycopersicum* | UniProt | Q588D6 |
| *Arabidopsis thaliana* | UniProt | Q9SR43 |
| *Posidonia australis* | OneKP | BYQM-2011005 |
| *Oryza sativa* | GenBank | BAD87875.1 |
| *Zea mays* | GenBank | NP_001105256.1 |

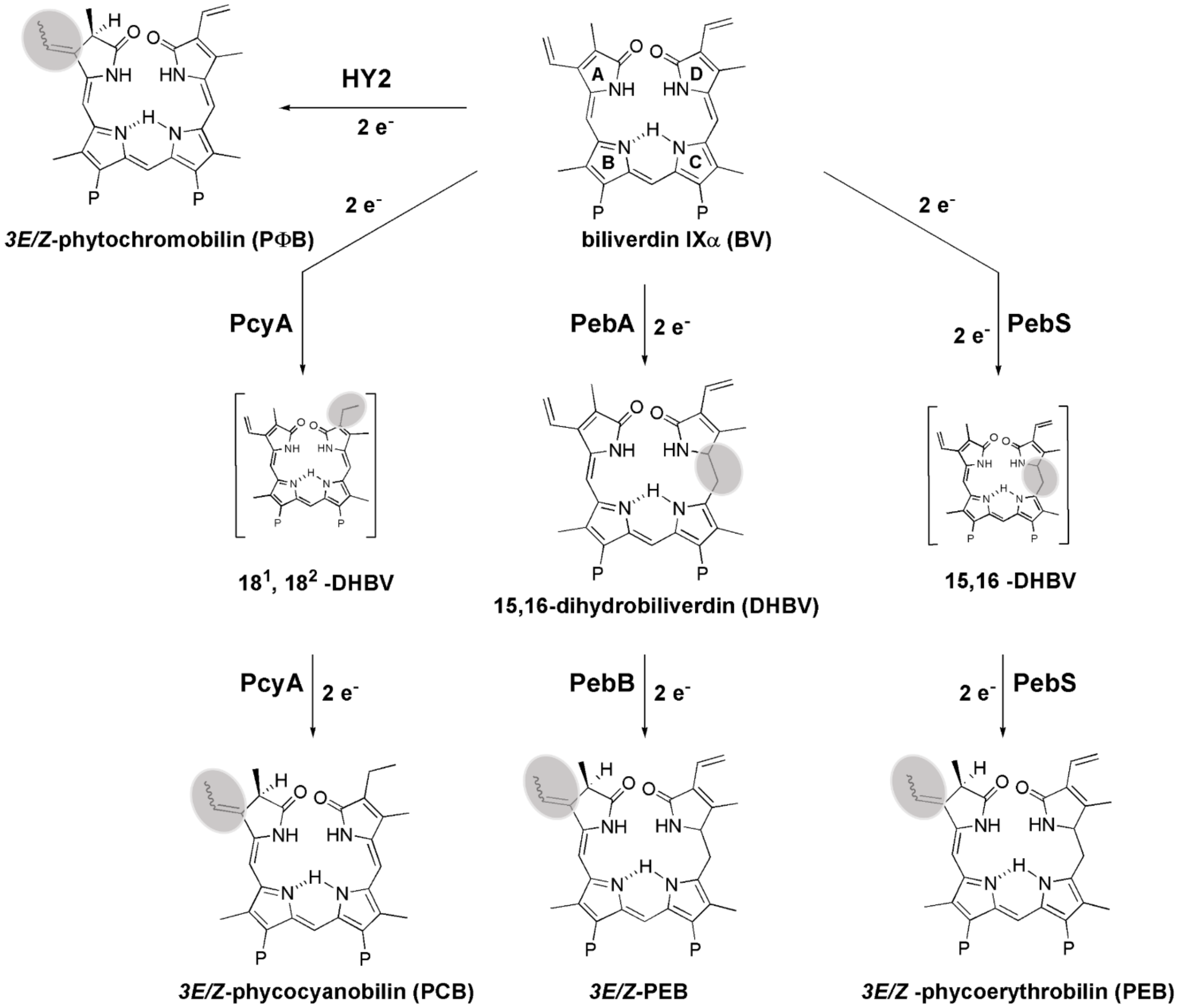


**Figure S1** FDBR family of bilin reductases. Reduction sites are highlighted in grey. The enzyme phytochromobilin synthase (HY2) is reducing BV at the A-ring to yield phytochromobilin (PΦB) (Frankenberg et al., 2001; Kohchi et al., 2001). PcyA is catalyzing the 4e$^-$ reduction of BV to phycocyanobilin (PCB) via the formation of $18^1$,$18^2$-dihydrobiliverdin ($18^1$,$18^2$-DHBV) as the intermediate. The enzymes PebA and PebB are working together to reduce BV to first 15,16-dyhrobiliverdin (15,16-DHBV) and ultimately to phycoerythrobilin (PEB). PebS combines the activity of PebA and PebB in one enzyme (Dammeyer et al., 2008). Thus far, HY2 has only been found in eukaryotic organisms whereas the other enzymes have been found in both prokaryotes and eukaryotes (PcyA, PebA, PebB) and PebS has only been detected in cyanophages.

```
                                                                                        105
                                                                                         ↓
PebS|P-SSM2                      ---------VTKDGGDAKL-----TANIRTGEHFLKAREAHIV-DP----NSDIYNTILY    92
PcyA|Synechocystis_PCC6803       LGY------VEGRLEGEKLV---IENRCYQTPQFRKMHLELAKVGK----GLDILHCVMF    92
PcyA|Prochlorococcus_MED4        LSN------IISNEEGKELY---IENEFYKAKGFRKLHIEVAEFSK----SLKILHCVFF    88
PcyA|Nostoc_PCC7120              LGY------VEGRLEGEKLT---IENRCYQTPQFRKMHLELAKVGN----MLDILHCVMF    89
PCYA|Chlamydomonas_reinhardtii   FSYADIPDPAKGEAGYPRLQ---LENRVYCSKVFRKLHVEVGLRQD----GLQVLHVVVY    279
PCYA|Cyanidioschyzon_merolae     LGY------IEGKLEGERVV---IENVCHQSPAFRKMHLELARVGTSQTASLDILHCVMF    154
PcyX|EBK42635                    -NLVEYNEPGMERFNNEKLG---WVNRTWNNRYIRRAHLDVVDVRE--SKGLWMAHLCLF    81
PebA|Synechococcus_WH8020        LEECRSS-----KSS---SV---IQSWLWDVPGFRRWRVTRLDAGD----SLQVFNSVAY    71
PEBA|Galdieria_sulphuraria       LASAQSS-----KKP---AR---IESWCYQCPKFRKIRLTYLDAGV----AAQVFNAVWY    143
PebB|Synechococcus_WH8020        FLQREDQ-----TGSKSKSIPVTTATWACKTEKFRQVRAACVSAGS----AASVLNFVIN    94
PEBB|Galdieria_sulphuraria       FEEKVAI-----TGKGSRQSTVKTTSYAYQSEKLRQIRAAHVQGGK----ALQVLNFVIF    141
PEBB|Guillardia_theta            YLRKEAM-----VGKGKRESLAWTQSYGYQTKKMRQIRAAHVNGGA----SLQVLNLVFF    88
HY2|Mesotaenium_caldariorum      YRCLQGM-----DGPRQKAV---SRSTALQSLKIRQMRSADISAGP----NLQVLNLVVF    78
HY2|Mesotaenium_endlicherianum   FRELIGD-----DGNGQVAV---SRSTAYQSEKLRQIRAADINAGP----FLQVLNLIAF    81
HY2|Interfilum_paradoxum         FIRLEAT-----DSKGQKAV---TDSTFLQSLKLRQVRAANIQAGN----NLQVLNLIMF    77
HY2|Cylindrocystis_brebissonii   FRLLRGM-----DGTCQQAL---SRSTALQSDKIRMFRSADISAGP----NLQVLNLVIF    85
HY2|Chlorokybus_atmophyticus     YGLLRAT-----DKKGDNAE---ARGDAFQTDKLRFIRPALITAGT----DFQVFNFVIM    75
KflaHY2                          FIRLEAT-----DSKGQKAV---TDSTYLQSHKLRQMRACNIQAGN----NLQVLNLIAF    109
HY2|Netrium_digitus              MRDMEGQ-----EGL-QGAI---CRTVAFQSDKIRLMRSADISAGK----NLQVLNLVIF    75
HY2|Ceratodon_purpureus          FRQMVAS-----DGQ---SL---IKNVAFESEKLRLFRCATINGGD----NMQVLNVAIC    96
HY2|Physcomitrella_patens        FQNMVAA-----DGQ---SV---VKNVAFESTKFRLFRCATINGGD----NMQVLNVAIC    191
HY2|Arabidopsis_thaliana         YSSMTGL-----DGK---TE---LQMLAFKSSKIRLLRSMAIEN-E----TMQVFDFAGF    120
HY2|Zea_mays                     FRFIKAN-----EDN---TV---LNALSFSTSKIRLLRSLTIEQKN----SVQVLDFAAF    83
HY2|Nicotiana_attenuata          YSSLLAM-----DDE---TE---LQMLSFEAPKIRLLRSLCIEGSD----GMQVLDFAAF    133
HY2|Solanum_lycopersicum         YSSLLSM-----DDK---TE---LQMLSFEAHKIRLLRSLCIEGTD----GMQVLDFAAF    127
                                                                  :   :               : .

                                              122
                                               ↓
PebS|P-SSM2                      PKT---GADLPCFGMDLMKFSDKK-VIIVFDFQHPREKY----------LFSVDGLP---    135
PcyA|Synechocystis_PCC6803       PEP---LYGLPLFGCDIVAGPGGV-SAAIADLSPTQSDR---Q-LPAAYQKSLAELG---    141
PcyA|Prochlorococcus_MED4        PDP---KYDIPIFGMDLVKVNELV-SAAIVDLSPSSKNQ---N-LK--YDHLLSHID---    135
PcyA|Nostoc_PCC7120              PRP---EYDLPMFGCDLVGGRGQI-SAAIADLSPVHLDR---T-LPESYNSALTSLN---    138
PCYA|Chlamydomonas_reinhardtii   PRY---SYDMPIFGMDIVMVDGRV-TLAVVDCCPVRADL---K-LQPHYMETMALLQRTF    331
PCYA|Cyanidioschyzon_merolae     PCAGRVGTNVPMFGCDIVAGRGAV-SAAIVDLSPVSASR---E-LPPSYRDALQSPEMAA    209
PcyX|EBK42635                    PML---TNGGPIYGFDIIAGEKKV-TGAFHDFSPLLQKD---HPLTKWFIEE----N---    127
PebA|Synechococcus_WH8020        PDY---NYDHPLMGVDLLWFGARQKLVAVLDFQPLVQDK---DYLDR-YFSGLKELN---    121
PEBA|Galdieria_sulphuraria       PIS---SLELPILGVDFLSFGGKN-VLCVMDFQPLMNDQ---YYLDK-YIQPLHPIR---    192
PebB|Synechococcus_WH8020        PKS---TYDLPFFGGDLVTLPAGH--LLALDLQPAIKTD---EVHTTHVWDRLIPIF---    143
PEBB|Galdieria_sulphuraria       PRL---EYDLPFFGADLVTLPGGH--LIALDMQPLFHTI---EYQNK-YSLQLQPIY---    189
PEBB|Guillardia_theta            PHM---NYDLPFLGLDLVTLPGGH--LIAIDMQPLFQTE---EYKKK-YAEPCMDMY---    136
HY2|Mesotaenium_caldariorum      PRP---EYDVPYFCADLVTISRGH--LVVLDLNPMYNTA---EYLDK-YITPLLPVH---    126
HY2|Mesotaenium_endlicherianum   PRP---EYDVPLLCADLVIFPRGQ--LLVLDLNPLFTTD---AYIDK-YIKPLLPRT---    129
HY2|Interfilum_paradoxum         PRP---EYDLPYFCADLVTLPRGH--LQIIDLNPLHGTQ---EHVEK-HIKPILPIT---    125
HY2|Cylindrocystis_brebissonii   PRP---AYDLPYFCADLVSIPRGH--LIVLDLNPLFCGE---EYVAK-HIRHILPLH---    133
HY2|Chlorokybus_atmophyticus     ARP---SFDLPIFGADFVCLPRGN--VIALDFQPLHQSP---SYLTK-YAERLLPLA---    123
KflaHY2                          PRP---EYDLPYFCADLVTLPRGH--LQILDLNPLHKTP---EHFEK-HIKPLLPIT---    157
HY2|Netrium_digitus              PKV---EYELPYFCADLVTFPRGH--LVVIDVNPMHNTD---VHFDR-FIRPFLPLR---    123
HY2|Ceratodon_purpureus          ARP---EYDLPIFCADFFSTPRMN--IVVLDLNPLYNTEQRPDYKEK-YFSPLLSMG---    147
HY2|Physcomitrella_patens        ARP---EYDLPIFCADFFSTARMN--IVVLDLNPLYNTEQRPEYKEK-YYSRILPLG---    242
HY2|Arabidopsis_thaliana         MEP---EYDTPIFCANFFTSTNVN--IVVLDLNPLHQLTDQTDYQDK-YYNKIMSIY---    171
HY2|Zea_mays                     SRP---EYDLPIFCANAFSSPARS--IVVLDLNPLYDTTEHKDYREK-YYRALMPLV---    134
HY2|Nicotiana_attenuata          PKP---EFDLPIFCANFFTAAKMN--IIVLDLNPLHDVMDQEDYKEK-YYKDLIPLG---    184
HY2|Solanum_lycopersicum         PKP---EFDLPIFCANFFTTAKMN--IIVLDLNPLHDIMDQEDYKEK-YYKDLITLG---    178
                                           *    : .            *

                                                                       242
                                                                        ↓
PebS|P-SSM2                      KEMIDNNKP-------VG------EDTTVYSDFDTYMTELDPVRGYMKNK-FGEGRSEAF    224
PcyA|Synechocystis_PCC6803       CHQSIVAEP--------LSEAQTLEHRQGQIHYCQQQQKNDKTRRVLEKA-FGEAWAERY    238
PcyA|Prochlorococcus_MED4        IQLSQSTSP-------DSDYEIIEERINYQKNYCVQQMKNEKTSLVLLKY-FDKVWVDEY    233
PcyA|Nostoc_PCC7120              CQGAIAASP--------VSAEQKQQILAGQHNYCSKQQQNDKTRRVLEKA-FGVDWAENY    235
PCYA|Chlamydomonas_reinhardtii   LTMSLNAVPVVAGPGDRREAARLQEIQDGQKRFCDNQLVNKKTRRVLEVA-MGVEWTEAY    438
PCYA|Cyanidioschyzon_merolae     CHLAQKSTS-------TPHLDQVREALEGQIHYCRKQQENDKTRRVLESA-FGKPWTERY    311
PcyX|EBK42635                    IDKIRN----------HEGEAEMADVIKAQNYYSEHQQKNPHTPRVMQSLGLPEEDIKLF    226
PebA|Synechococcus_WH8020        WDLHDNAKS-------IPSTIPPEEVKNLQDKYDIYSAERDPAHGLFTSH-FGKDWSNRF    223
PEBA|Galdieria_sulphuraria       ITLAKQ--T-------VEDDKSRSIIWQLQCEYDRYSAEKDPAMSLFRSY-FGSTWAHRF    293
PebB|Synechococcus_WH8020        LELAASAER-------VTDERSE-VLLQGQRKYTDYRAEKDPARGMLTRF-HGSEWTEAY    249
PEBB|Galdieria_sulphuraria       LKFVNHAVH-------IQDKQLLELIKSRHLAYIQYRAEKDPARGMFTRF-YGSEWTEEY    292
PEBB|Guillardia_theta            LDFVEAAKP-------VTDPDHLARIRERQLSYLQYRAEKDPARGMFTRM-YGPEWTERY    239
HY2|Mesotaenium_caldariorum      LGMLNSAVA-------ETDEMKILSNKLAQHRYMSWRCEKDPGRPVLTRL-FGSERCEEY    229
HY2|Mesotaenium_endlicherianum   IDMLEAAQP-------ETDPDQIAKNQEAQHRYMSWRAEKDPGRPLLTRL-FGEERCERF    234
HY2|Interfilum_paradoxum         ADMVAKAEP-------TSDPIEIARNREAQHKYISWRAEKDPGRPVITRL-YGTELCEAY    228
HY2|Cylindrocystis_brebissonii   LDMLDAAEP-------ATDPAVIAANQEAQHRYICWRCEKDPGRPVLTRL-FGEKRCESY    236
HY2|Chlorokybus_atmophyticus     LDYVDSAEA-------TTDSRAEEANRDAQHKYLTYRAVKDPGRGVLTRY-FGAELTEAY    226
KflaHY2                          AEMVAKAVP-------TTDPVEIARNREAQHKYVCWRAEKDPGRPTITRL-YGTELCEAY    260
HY2|Netrium_digitus              LAMLEECQP-------ERDPSRLAANAEAQHRYMCWRAEKDPGRPVLTRL-FGTERCEKL    226
HY2|Ceratodon_purpureus          LDMAEKAEP-------SEDADEVAENRESHHRYLMWRATKDPGRYILMRL-YGEELCERY    250
HY2|Physcomitrella_patens        LDMADKANP-------SNDAYEIAENQESHHRYLMWRATKDPGRYLLMRL-FGEPLCERY    345
HY2|Arabidopsis_thaliana         LEMTIQVRE-------EMEPSHVRANCEAQHKYLTWRAQKDPGHGLLKRL-VGEAKAKEL    274
HY2|Zea_mays                     LEFMDGAVR-------ESSKEKIDRNREAQHKYLTWRAEKDPGYPLLKKL-IGESGAKDL    237
HY2|Nicotiana_attenuata          LELTDKSEE-------ETDASQVARNREAQHRYLTWRSEKDPGHGVLKRL-VGEALAKDV    287
HY2|Solanum_lycopersicum         LGLMDRSEG-------ETDASQIACNCEAQHRYLTWRSEKDPGHGVLKRL-IGEDLAKDV    281
                                                                 :             :          .
```

**Figure S2** FDBRs sequence alignment. The alignment was constructed using ClustalW (https://www.ebi.ac.uk/Tools/msa/clustalo/). The amino acid residues mentioned in the text are highlighted in different colors and labeled according to KflaHY2. PebS|P-SSM2: Q58MU6 (UniProt). PcyA|Synechocystis_PCC6803: Q55891 (UniProt). PcyA|Prochlorococcus_MED4: Q93TL5. PcyA|Nostoc_PCC7120: Q93TN0 (UniProt). PCYA|Chlamydomonas_reinhardtii: A8HUP3 (UniProt). PCYA|Cyanidioschyzon_merolae: CYME_CMH110C (Genbank). PcyX|EBK42635: GOS_8734801. PebA|Synechococcus_WH8020: Q02189 (UniProt). PEBA|Galdieria_sulphuraria: M2XA99 (UniProt). PebB|Synechococcus_WH8020: Q02190 (UniProt). PEBB|Galdieria_sulphuraria: M2X9Q0 (UniProt). PEBB|Guillardia_theta: AIA66937.1 (UniProt). HY2|Mesotaenium_caldariorum: HKZW-2005125 (OneKP). HY2|Mesotaenium_endlicherianum: WDCW-2047152 (OneKP). HY2|Interfilum_paradoxum: FPCO-2003903 (OneKP). HY2|Cylindrocystis_brebissonii: YOXI-2055982 (OneKP). HY2|Chlorokybus_atmophyticus: AZZW-2019954. KflaHY2: GAQ84116.1 (GenBank). HY2|Netrium_digitus: FFGR-2000050 (OneKP). HY2|Ceratodon_purpureus: FFPD-2057880 (OneKP). HY2|Arabidopsis_thaliana: Q95R43 (UniProt). HY2|Zea_mays: Q6E5A3 (UniProt). HY2|Nicotiana_attenuata: A0A1J6J6Q1 (UniProt). HY2|Solanum_lycopersicum: Q588D6 (UniProt).

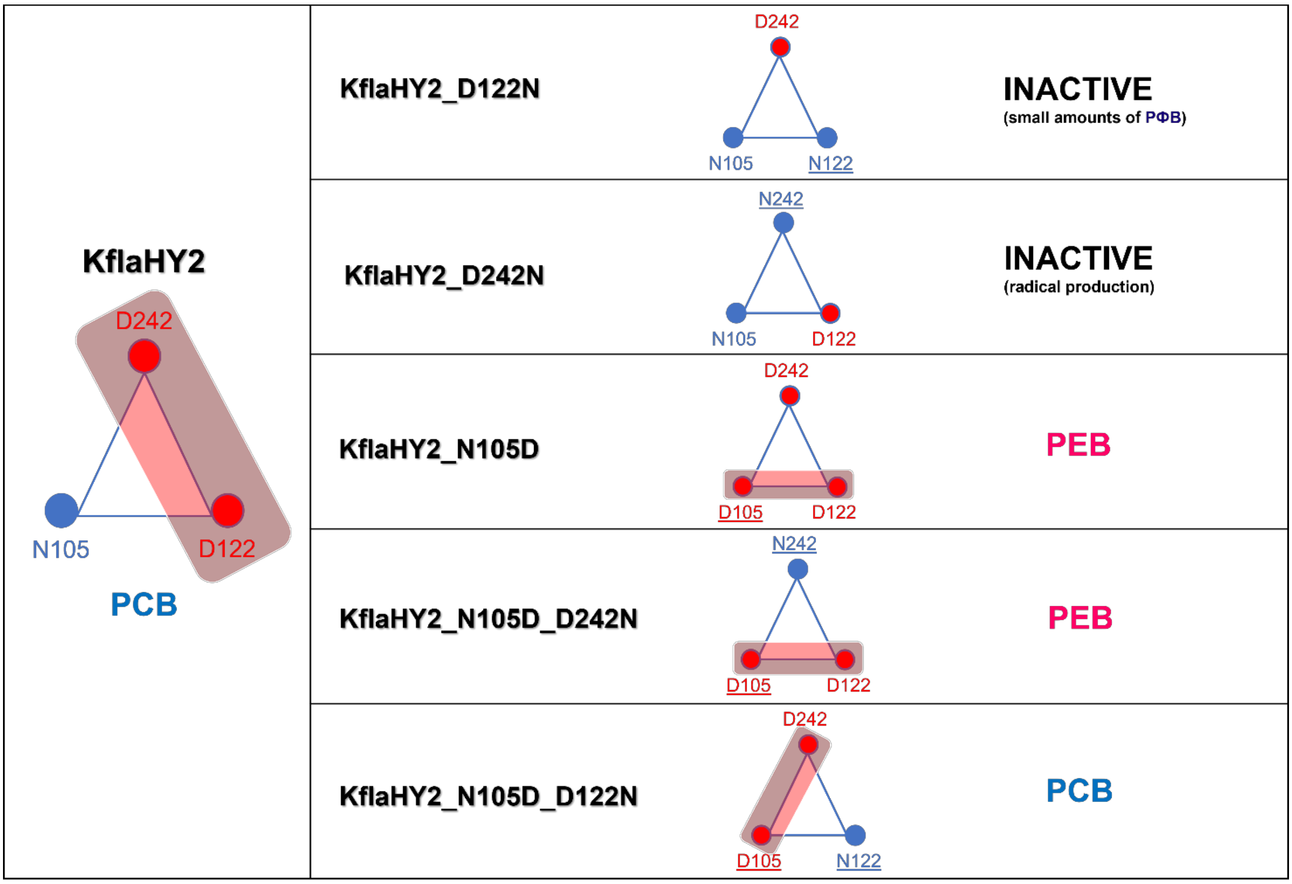

**Figure S3** Schematical representation of the active sites and activity of KflaHY2 and its variants. The red box highlights the position of the two aspartate residues. Exchanged residues are underlined. The production of either PCB or PEB is determined by the position of aspartate couple in the active site. Variants in which the aspartate residues are opposite to each other are catalyzing BV reduction to PCB. Variants in which the aspartates are laying on the same side show reductases-isomerase activity, first catalyzing BV reduction to PCB and then isomerizing it to PEB.
